# Spt4 drives cellular senescence by activating non-coding RNA transcription in ribosomal RNA gene clusters

**DOI:** 10.1101/2022.04.22.488906

**Authors:** Masaaki Yokoyama, Mariko Sasaki, Takehiko Kobayashi

## Abstract

Genome instability can drive aging in many organisms. The ribosomal RNA gene (rDNA) cluster is one of the most unstable regions in the genome. Replicative lifespan in budding yeast is correlated to rDNA stability, suggesting that the rDNA locus produces an aging signal. To understand the underlying mechanism, we looked for yeast mutants with more stable rDNA and longer lifespan than wild-type cells. We reveal that absence of a transcription elongation factor, Spt4, resulted in an increased rDNA stability, a reduced activity of the regulatory E-pro promoter in the rDNA, and extended replicative lifespan in a *SIR2*-dependent manner. Spt4-dependent lifespan restriction was abolished in the absence of non-coding RNA transcription at the E-pro locus. The amount of Spt4 increases and its function becomes more important as cells age. These findings suggest that Spt4 is a promising aging factor that accelerates cellular senescence through rDNA instability driven by non-coding RNA transcription

## Introduction

Aging is the decline of the biological functions, which occurs over time and involves complex biological processes (López-Otín et al., 2013; McHugh and Gil, 2018; Di Micco et al., 2021). Aging is the critical risk factor that trigger the onset of many diseases such as cancer, neurological diseases, and cardiovascular diseases. Therefore, understanding the molecular mechanisms of aging is important for designing therapies to delay or prevent aging-related diseases. One of the causes of aging is cellular senescence (López-Otín *et al*., 2013; McHugh and Gil, 2018; Di Micco *et al*., 2021). Budding yeast, *Saccharomyces cerevisiae*, where genetic manipulation experiments can be easily done, has been used as a model organism to elucidate the molecular mechanisms underlying cellular senescence (Sinclair et al., 1998b; Longo et al., 2012; He et al., 2018).

The rDNA encodes ribosomal RNA transcription unit(s) and is one of the highly abundant sequences that are organized into array(s) of tandem repeats at a single or several loci in eukaryotic genomes (reviewed in (Kobayashi, 2011)). Due to this repetitive arrangement, the rDNA can change in copy number. The budding yeast genome contains ∼150 rDNA copies at a single locus on chr XII (Fig. 1A). Each copy contains 35S and 5S rRNA transcription units that are separated by two intergenic spacers (IGSs). The IGS1 contains a replication fork barrier (RFB) site and an RNA polymerase (RNAP) II-dependent, bi-directional promoter, E-pro, that synthesizes non-coding RNAs and the IGS2 contains an origin of DNA replication and cohesin-associated region (Fig. 1A). After DNA replication is initiated, the replication fork moving in the direction opposite to the 35S rDNA is arrested at the RFB site bound by a Fob1 protein, leading to formation of DNA double-strand breaks (DSBs) (Brewer et al., 1992; Kobayashi et al., 1992; Kobayashi and Horiuchi, 1996; Weitao et al., 2003; Burkhalter and Sogo, 2004; Kobayashi et al., 2004). Previous studies have identified two major processes that dictate the outcomes of the DSB repair as to whether it results in rDNA copy number changes and thus rDNA instability. First, end resection of DSBs, an initiating event for homologous recombination (HR), is normally suppressed at the RFB (Sasaki and Kobayashi, 2017). Thus, removal of this suppression can lead to HR-mediated repair (Fig. 1A) (Sasaki and Kobayashi, 2017). If DSB repair engages the rDNA copy at an aligned position on the sister chromatid, it does not lead to a change in the rDNA copy number (Kobayashi, 2011). However, if DSB repair involves a misaligned copy on the sister chromatid or another copy on the same chromosome, expansion or contraction of the rDNA array will be the outcome. Furthermore, such DSB repair can lead to the production of extrachromosomal rDNA circles (ERCs). Second, transcription of non-coding RNA from E-pro is normally repressed by a histone deacetylase Sir2, which results in stable association of cohesin to the rDNA (Fig. 1A) (Kobayashi *et al*., 2004; Kobayashi and Ganley, 2005). This repression is relieved by the absence of Sir2 or in a strain with a low number of rDNA units where *SIR2* expression is reduced, leading to enhanced transcription from E-pro, disruption of cohesin association and changes in rDNA copy number (Kobayashi *et al*., 2004; Kobayashi and Ganley, 2005; Iida and Kobayashi, 2019).

**Figure 1.**
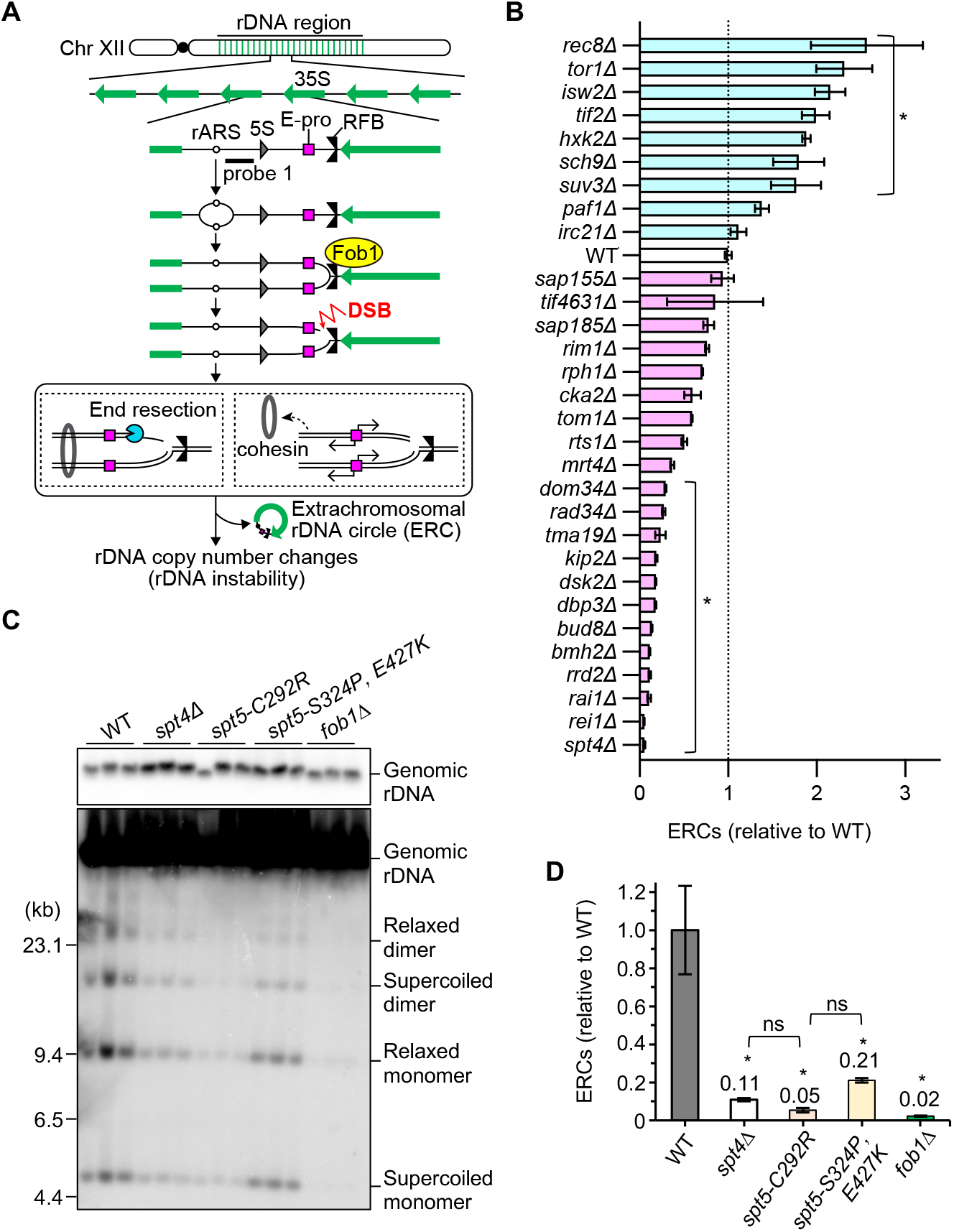
The Spt4-Spt5 complex enhances rDNA instability by promoting ERC production. **(A)** DNA replication fork arrest, DSB formation and copy number changes in the budding yeast rDNA region. 35S, 35S precursor rRNA coding region; 5S, 5S rRNA coding region; rARS, autonomously replicating sequence in rDNA; RFB, replication fork barrier; DSB, DNA double-strand break; ERC, extrachromosomal rDNA circle. The black bar indicates the position of probe 1 used for ERC detection by Southern blotting. **(B)** The summary of the ERC analyses. See Figure S1 for the raw data. The level of ERCs for the indicated strains was determined by calculating the ratio of ERCs relative to genomic rDNA and normalized to the average level of ERCs in wild-type clones (WT; bar shows mean ± s.e.m; bars for mutant strains show the range between the ERC-levels for two independent clones). One-way ANOVA was performed for multiple comparisons (asterisks indicate statistically significant difference between wild type and mutants [*p* < 0.05]). **(C)** ERC detection. DNA was isolated from three independent clones of the indicated strains and separated by agarose gel electrophoresis, followed by Southern blotting with probe 1 shown in (A). Genomic rDNA, supercoiled and relaxed forms of monomeric and dimeric ERCs are indicated. Sizes of lambda DNA-*Hind* III markers are indicated. **(D)** Quantitation of ERC levels. ERCs in (C) were quantified, as described in (B). Bars show mean ± s.e.m. One-way ANOVA was performed for multiple comparisons (asterisks indicate statistically significant difference between wild type and mutants [*p* < 0.05]; ns indicates no significant difference (*p* > 0.05)).

The budding yeast cells produce a finite number of daughter cells before death, which defines the replicative lifespan (Sinclair et al., 1998a). The replicative lifespan of cells lacking *SIR2* is shortened to half of that of the wild-type strain, while it is extended by over-production of this protein (Kaeberlein et al., 1999). The amount of this protein decreases in old cells (Fine et al., 2019). Therefore, Sir2 works as an anti-aging factor. On the contrary, in the *fob1* mutant, the rDNA is stable and the lifespan is extended by ∼60% compared to the wild-type strain (Kobayashi et al., 1998; Takeuchi et al., 2003). In addition, a strain in which the E-pro is replaced with a galactose-inducible bi-directional *GAL1/10* promoter, hereafter referred to as the Gal-pro strain, has stable rDNA and longer lifespan in glucose medium while the opposite is observed when transcription is induced by the addition of galactose in the medium (Kobayashi and Ganley, 2005; Saka et al., 2013). These findings demonstrate that rDNA stability can dictate replicative lifespan.

To better understand how cellular senescence can be regulated via the rDNA locus, we aim to identify factors that are involved in the control of both processes. The presence of anti-aging factors such as Sir2 inspired us to find proteins with the opposite function, which contribute to aging by destabilizing the rDNA cluster and would be present in increased amounts or activated in old cells. To identify such aging factors, we screened for long-lived yeast mutants with stable rDNA and found that cells lacking *SPT4* showed increased rDNA stability and extended lifespan.

The progression of the RNAP II transcription machineries can be paused by various factors, including the nucleosomes that are intrinsic barriers to RNAP II progression (Ehara and Sekine, 2018; Kujirai and Kurumizaka, 2020). To ensure efficient transcription, organisms have evolved factors that enhance the processivity of RNAP II. Spt4 forms a heterodimer with Spt5, known as the Spt4-Spt5 complex (and, in metazoans, as DSIF, the DRB sensitivity-inducing factor - DRB is 5,6-dichloro-1-beta-d-ribofuranosyl-benzimidazole). The Spt4-Spt5 complex interacts with RNAP II and plays an important role in facilitating transcription by RNA polymerase II in the context of chromatin structures (Hartzog and Fu, 2013; Ehara and Sekine, 2018; Kujirai and Kurumizaka, 2020; Decker, 2021). The Spt4-Spt5 complex also interacts with RNAP I and localizes to the rDNA, while the influence of this complex on rRNA transcription can be both positive and negative (Schneider et al., 2006; Anderson et al., 2011; Lepore and Lafontaine, 2011; Viktorovskaya et al., 2011). In cells lacking Spt4, transcription of non-coding RNA from E-pro was reduced, while rDNA stability and replicative lifespan increased in a manner dependent on *SIR2*, *FOB1*, and E-pro activity. Moreover, in old cells, an increase in the amount of Spt4 was observed, which possibly enhances the impact of the protein on facilitating transcription from E-pro as cells age, accelerating cellular senescence. Our results suggest that Spt4 is a promising aging factor that drives cellular senescence through destabilizing rDNA in budding yeast.

## Results

### The Spt4-Spt5 complex enhances rDNA instability and shortens the replicative lifespan

In this study, we sought to identify the putative factors that spread the aging signal and restrict the replicative lifespan by destabilizing rDNA. Cells lacking such factors would display increased lifespan and rDNA stability. A previous study has determined the replicative lifespan of 4,698 mutant strains that lack non-essential genes for viability and identified 238 mutant strains with increased lifespan (McCormick et al., 2015). Among them, 30 mutants lack a gene that has an annotated function in DNA metabolism, such as recombination, replication, and repair. We examined the degree of rDNA stability in these mutants by measuring the amount of ERCs (Fig. 1A), which are often produced during rDNA copy number changes and can be used as an indicator of rDNA instability (Sinclair and Guarente, 1997; Kaeberlein *et al*., 1999; Ganley et al., 2009). Measurements of ERCs identified 7 mutants with, compared to wild-type cells, a statistically significant increase in ERCs and 12 mutants with decreased ERC levels, the latter of which displayed the expected phenotype for cells that lack aging factors of our interest (Fig. 1B, Fig. S1A–D).

Cells lacking Spt4 accumulated the least amount of ERCs among the mutants examined (Fig. 1B). Spt4 forms a heterodimer with Spt5, known as the Spt4-Spt5 complex and functions as a transcription elongation factor that facilitates transcription by RNA polymerase II (Hartzog and Fu, 2013; Ehara and Sekine, 2018; Kujirai and Kurumizaka, 2020; Decker, 2021). Spt4 also facilitates rRNA transcription by RNA polymerase I and is localized across the rDNA region (Schneider *et al*., 2006; Anderson *et al*., 2011; Lepore and Lafontaine, 2011). Our study revealed that Spt4 has a novel function destabilizing rDNA. We sought to understand how this protein reduces rDNA stability and induces cellular senescence.

To determine whether Spt4 enhances rDNA instability alone or through the Spt4-Spt5 complex, we tested the involvement of Spt5 in the regulation of rDNA stability. As *SPT5* is an essential gene, two temperature sensitive alleles, *spt5-*S324P, E427K and *spt5-*C292R, were used (Anderson et al. 2011). Serine 324, substituted in the first mutant, is located within the conserved NusG-like domain of Spt5 and is required for binding to Spt4 (Guo et al., 2008). Thus, the *spt5-*S324P, E427K mutant is deficient with the Spt4-Spt5 complex formation. A point mutation in the second allele results in substitution of cysteine 292, which, however, does not lie on the interface with Spt4 (Anderson et al. 2011). Because the *spt5-*C292R mutant shows a reduction in the synthesis rate of rRNA, this allele compromises the function of Spt5 in the Spt4-Spt5 complex (Anderson et al. 2011). These strains were grown at 27°C and analyzed for ERCs. The level of ERCs in the *spt4*Δ mutant was ∼10-fold lower than that in wild-type cells but this level was still ∼5-fold higher than that in the mutant lacking *FOB1* (Fig. 1C, 1D), which is responsible for DNA replication fork arrest and ERC production (Kobayashi and Horiuchi, 1996; Defossez et al., 1999). The two *spt5* mutants showed a statistically significant reduction in the level of ERCs, compared to wild type (Fig. 1C, 1D). The ERC levels in the *spt5-*C292R and *spt5-*S324P, E427K mutants were ∼2-fold lower and ∼2-fold higher than in the *spt4*Δ mutant, respectively, but these mutual differences were not statistically significant. Overall, these results demonstrate that Spt4 and Spt5 function as a complex when inducing rDNA instability.

### Spt4-mediated rDNA destabilization depends on *SIR2*

We examined whether reduced ERC formation paralleled an increase in rDNA stability in the *spt4*Δ mutant by assessing the degree of size heterogeneity of chr XII by pulsed-field gel electrophoresis (PFGE). The band of chr XII in the *spt4*Δ mutant appeared as sharp as that in the *fob1*Δ mutant defective in rDNA copy number changes and seemed sharper than that in wild-type cells in the *SIR2* background (Fig. 2A). A previous study reported that the average rDNA copy number is reduced in absence of Spt4 by ∼3 fold, compared to wild-type cells (Schneider *et al*., 2006). We observed some variablities in the size of chr XII among four independent *spt4*Δ clones constructed for our study, but not matching a 3-fold change in the case of samples with increased or decreased rDNA copy number (Fig. 2A, WT vs. *spt4*Δ in the *SIR2* background).

**Figure 2.**
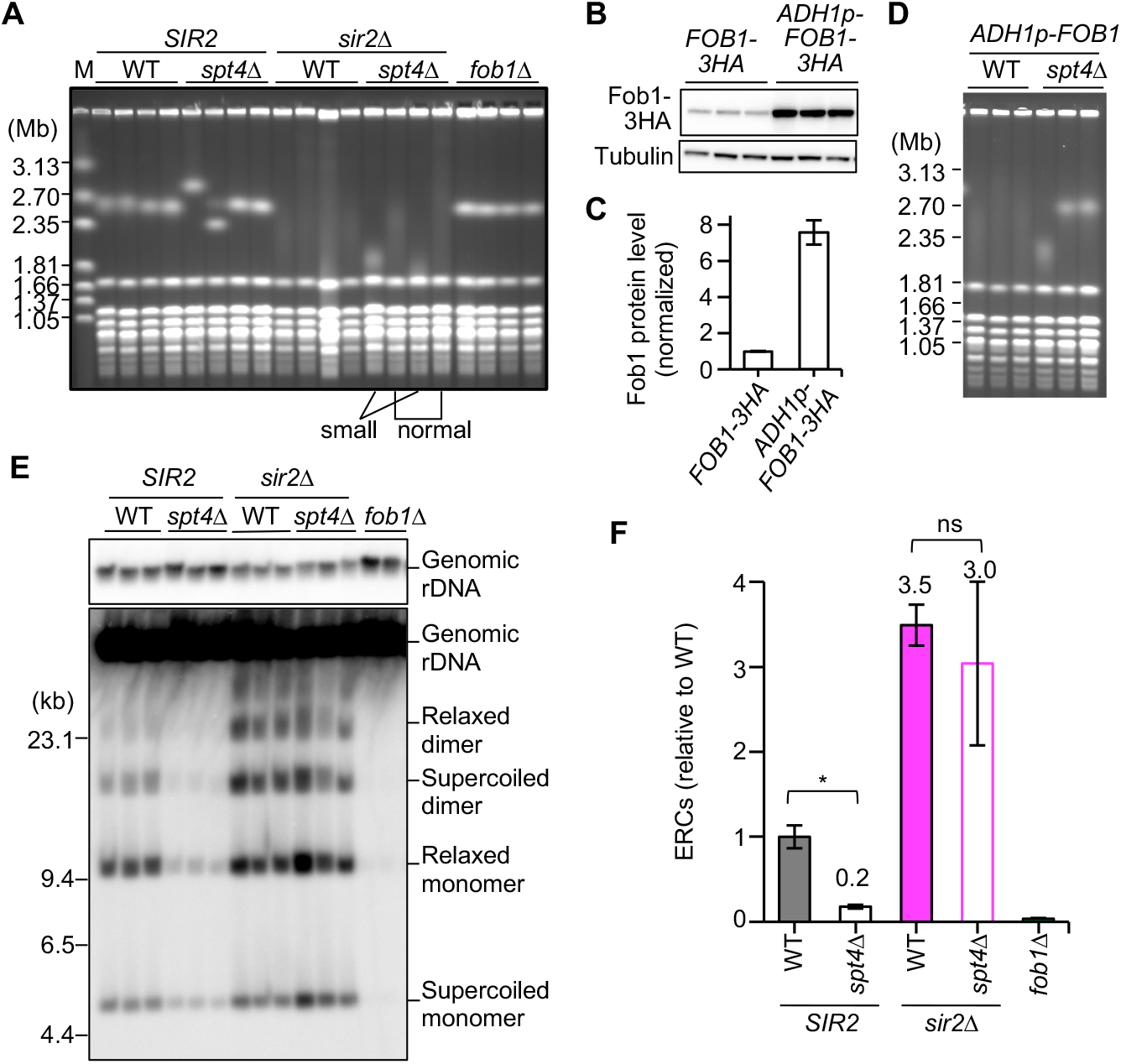
Spt4-mediated enhancement of rDNA instability requires *SIR2*. **(A, D)** Size heterogeneity of chromosome XII. DNA was extracted from four (A) and three (D) independent clones of the indicated strains and separated by PFGE. DNA was stained with ethidium bromide (EtBr). The extent of variability in rDNA copy number determines the appearance of the chr XII band; stable rDNA migrates together, while a diffused appearance of the band (from little to extreme smearing) reflects (small to large) differences in rDNA copy number in cell population. In (A), four independent clones of the *sir2*Δ *spt4*Δ mutant were marked by either ’small’ or ’normal’, depending on the size of chr XII. M indicates *Hansenula wingei* chromosomal DNA markers. **(B)** Overproduction of Fob1 protein. Fob1, C-terminally tagged with a triple HA epitope (Fob1-3HA), was detected in protein extracts prepared from three independent clones of the indicated strains that carry the *FOB1* ORF under control of either its endogenous promoter (*FOB1-3HA*) or the constitutive *ADH1* promoter (*ADH1p-FOB1-3HA*). Proteins were separated by SDS-PAGE, followed by Western blotting with antibodies against the HA tag and tubulin. **(C)** Quantitation of Fob1 protein levels. The levels of Fob1-3HA in (B) were quantified relative to tubulin, which was normalized to the average of *FOB1-3HA* clones (bars show mean ± s.e.m). **(E)** ERC detection. DNA isolated from the indicated strains was separated by agarose gel electrophoresis, followed by Southern blotting with probe 1 (shown in Figs. 1A and 3A). Genomic rDNA, supercoiled and relaxed forms of monomeric and dimeric ERCs are indicated. Sizes of lambda DNA-Hind III markers are shown. **(F)** Quantitation of ERC levels. Levels of ERCs in (E) were determined for the indicated strains as the ratio of the sum of monomeric and dimeric ERCs relative to genomic rDNA (top panel), which was normalized to the average of wild-type clones (bars show mean ± s.e.m). Statistical analyses were performed between wild type and *spt4*Δ and between *sir2*Δ and *sir2*Δ *spt4*Δ, by two-sided Welch t test. A significant difference (*p* < 0.05) is marked by an asterisk; ns indicates no significant difference (*p* > 0.05).

To determine more precisely the effect of *SPT4* deletion on rDNA stability, we overexpressed Fob1 by placing the *FOB1* ORF under the control of the constitutive *ADH1* promoter, which led to ∼8-fold increase in Fob1 protein level (Fig. 2B, 2C). Compared to the wild type (Fig. 2A, lanes WT/*SIR2)*, Fob1 overexpression resulted in smearing of the chr XII band (Fig. 2D, lanes WT/*ADH1p-FOB1*). Upon deletion of *SPT4* from the *ADH1p-FOB1* strain, the chr XII band became far less smeared (Fig. 2D). Therefore, *SPT4* induces copy number changes in the chromosomal rDNA array, and thus rDNA instability.

The histone deacetylase Sir2 is a key player in the suppression of both rDNA instability and cellular senescence (Kaeberlein *et al*., 1999; Kobayashi *et al*., 2004), thus with a role opposite to that of Spt4. To gain insight into the relationship between *SPT4* and *SIR2* during processes that affect rDNA stability, we constructed a double mutant of *spt4*Δ and *sir2*Δ and examined rDNA stability (Fig. 2A, 2E, 2F). In the PFGE assay, the chr XII band of *sir2*Δ cells was extremely smeared (Fig. 2A, lanes *WT/sir2*Δ), consistent with a previous study (Kobayashi *et al*., 2004). The chr XII band in two of the four independent *sir2*Δ *spt4*Δ clones examined, was as smeared as that in the *sir2*Δ clones (Fig. 2A, lanes marked by ’normal’ vs. WT/*sir2*Δ), while the other two clones showed sharper bands but carried a much shorter chr XII (Fig. 2A, lanes marked by ’small’). It remains unclear whether the reduction in the rDNA cluster size is a direct effect of constructing the *SPT4* deletion in the *sir2*Δ strain background. It is known that transformation procedures that are used for gene replacement can induce rDNA copy number changes (Kwan et al., 2016). Thus, it is likely that these two clones have spontaneously lost rDNA copies during strain construction. Because chr XII in these clones migrated much faster than the *sir2*Δ and other two *sir2*Δ *spt4*Δ clones (Fig. 2A), it was difficult to compare the size heterogeneity of chr XII band because resolution of DNA molecules is expected to be very different between two regions. Thus, it appears that removal of Spt4 protein does not counteract rDNA instability caused by *SIR2* deletion.

Deletion of *SPT4* from the WT (*SIR2*) cells resulted in a decrease in ERCs by ∼5-fold (Fig. 2E, 2F). However, the ERC level in the *spt4*Δ *sir2*Δ double mutant was similar or only slightly reduced, compared to the *sir2*Δ single mutant (Fig. 2F, WT vs. *spt4*Δ in the *sir2*Δ background), consistent with the outcome of the PFGE analysis (Fig. 2A). Taken together, these findings indicate that Spt4-mediated enhancement of rDNA instability depends on *SIR2*. In other words, Spt4 functions downstream of Sir2 in the regulation of rDNA stability.

### Spt4 enhances transcription of non-coding RNA from E-pro

Previous studies have identified several factors that impact rDNA stability. Fob1-dependent replication fork arrest at the RFB site leads to formation of DSBs (Weitao *et al*., 2003; Burkhalter and Sogo, 2004; Kobayashi *et al*., 2004) (Fig. 1A). When these DSBs are resected, HR-dependent repair is taken place that can induce rDNA copy number changes (Sasaki and Kobayashi, 2017). We first examined the frequency of replication fork arrest by two-dimensional (2D) agarose gel electrophoresis. Genomic DNA was isolated from WT and *spt4*Δ mutant cells and digested with the restriction enzyme *Nhe* I (Fig. 3A). DNA fragments were separated by agarose gel electrophoresis according to molecular mass in the first dimension and then according to molecular mass and shape in the second dimension, followed by Southern blotting. The level of arrested forks in total replication intermediates was similar between WT and *spt4*Δ strains, thereby indicating that RFB activity is not reduced in the *spt4*Δ mutant (Fig. 3B, 3C).

**Figure 3.**
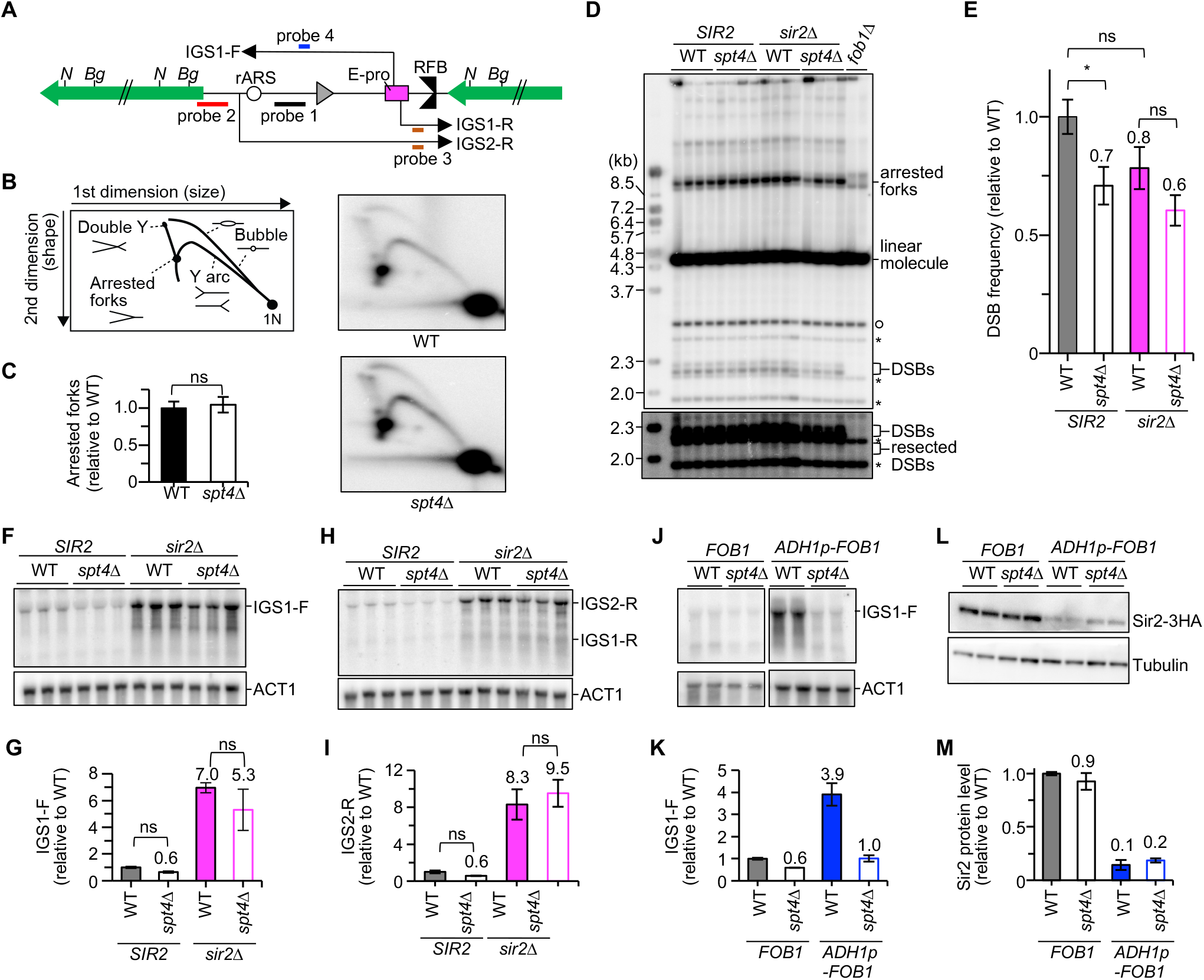
Spt4 stimulates transcription of non-coding RNA from E-pro. **(A)** Restriction map and the positions of the probes used in this study. *N* and *Bg* indicate restriction sites recognized by *Nhe* I and *Bgl* II, respectively. The 35S (green) and 5S (gray) rRNA genes (arrow bars), the origin of replication within the rDNA (rARS) and the regulatory, bidirectional promoter E-pro are shown. Positions of probes used for 2D (Probe 1) and DSB (probe 2) analyses in (B) and (D) are indicated, as well as the position of single-stranded probes used for detection of non-coding RNA transcribed from E-pro in the direction toward the RFB site (Probe 3) or the 5S gene (Probe 4). **(B)** 2D agarose gel electrophoresis. Genomic DNA isolated from the indicated strains was digested with *Nhe* I. DNA was separated by size in the first dimension and by size and shape in the second dimension, followed by Southern blotting with probe 1, as indicated in (A). The diagram on the left shows the expected migration pattern of different replication intermediates. 1N represents the bulk of unreplicated, linear DNA. **(C)** Quantitation of fork arrest. The signals of arrested forks in (B) were quantified relative to the total of replication-intermediate signals (that included bubble arcs, Y arcs, arrested forks, and double Y spots). The levels of arrested forks in the *spt4*Δ mutant was normalized to the average of wild-type clones. **(D)** DSB assay. Genomic DNA isolated from the indicated strains was digested with *Bgl* II. DNA was separated by single-dimension agarose gel electrophoresis, followed by Southern blotting with probe 2 indicated in (A). Intermediates are indicated incl. terminal fragments containing the telomere-proximal rDNA repeat (open circle) and non-specific background (asterisks). In the absence of Fob1 (*fob1*Δ), intermediates with replication forks that just entered the 35S rRNA region are prevalent. **(E)** Quantitation of DSBs. The DSB frequency was determined by quantifying the signal of DSBs in (D) relative to that of arrested forks, which was normalized to the average of wild-type clones (bars show mean ± s.e.m). One-way ANOVA was performed for multiple comparisons. An asterisk indicates a significant difference (*p* < 0.05); ns marks a difference that is not statistically significant (*p* > 0.05). **(F, H, J)** Detection of non-coding RNAs transcribed from E-pro. RNA isolated from the indicated strains was separated by gel electrophoresis in formaldehyde-containing agarose gel, followed by Northern blotting. IGS1-F transcribed from E-pro toward 5S rDNA was detected with probe 4 (F, J). RNA transcribed toward the RFB from E-pro (IGS1-R) or from an upstream promoter (IGS2-R) were detected with probe 3 (H). Membranes were reprobed for *ACT1* transcripts as a control for loading (ACT1). **(G, I, K)** Quantitation of non-coding RNA. Signals corresponding to IGS1-F (G, K) and IGS2-R transcripts (I) were quantified relative to ACT1, which was normalized to the average of wild-type clones. Bars show mean ± s.e.m. in (G) and (I) and mean and the range from two independent experiments in (K). Signals of IGS1-R transcripts were detected below detection limit and were thus not quantified. One-way ANOVA was performed for multiple comparisons in (G) and (I). An asterisk indicates a significant difference (*p* < 0.05); ns marks a difference that is not statistically significant (*p* > 0.05). **(L)** Detection of Sir2 that is C-terminally tagged with a triple HA epitope (Sir2-3HA). Protein extracts isolated from four independent clones of the indicated strains were separated by SDS-PAGE, followed by Western blotting with antibodies against HA tag and tubulin. **(M)** Quantitation of Sir2 protein levels. The level of Sir2-3HA protein was quantified relative to tubulin, which was normalized to the average of wild-type clones (bars show the range from two independent experiments).

We also determined the DSB frequency at the RFB site by separating genomic DNA digested with the restriction enzyme *Bgl* II by conventional agarose gel electrophoresis and subsequent Southern blotting. The DSB frequency was reduced in the *spt4*Δ mutant by ∼30%, compared to WT cells (Fig. 3D, 3E). Deletion of *SPT4* also caused a reduction in DSB frequency in the *sir2*Δ background but the difference was not statistically significant (Fig. 3D, 3E). We did not use cells synchronized in S phase for DSB assays, which made assessing the frequency of resected DSBs unreliable as resected DSB intermediates were mostly below the detection limit of Southern blotting (Fig. 3D, lower panel). Thus, the reduction of DSBs and possible reduction of DSB end resection might be reflected in the increased rDNA stability in the *spt4*Δ mutant, which needs to be determined in future studies.

Sir2 represses transcription of non-coding RNA from E-pro when the copy number is normal (Kobayashi and Ganley, 2005; Iida and Kobayashi, 2019). Relief of this repression activates transcription from E-pro, which induces disruption of cohesin binding and rDNA instability during DSB repair (Fig. 1A) (Kobayashi *et al*., 2004; Kobayashi and Ganley, 2005; Saka *et al*., 2013). Chromatin immunoprecipitation experiments indicate that Spt4 is localized across the rDNA, being enriched around the middle of the 35S rDNA gene and IGS1 (Lepore and Lafontaine, 2011). Spt5 is also localized across the rDNA but, interestingly, is mostly enriched at the IGS1 near the RFB and E-pro (Lepore and Lafontaine, 2011). This raises the possibility that Spt4, as part of the Spt4-Spt5 complex functions to destabilize the rDNA through transcription regulation of non-coding RNA from E-pro. To test this possibility, we detected the noncoding RNA transcripts by Northern blotting with strand-specific probes (Fig. 3A). There are three major non-coding RNA species that are transcribed in intergenic spacer regions: IGS1-F and IGS1-R are transcribed from E-pro toward the 5S rRNA gene and the RFB site, respectively (Kobayashi and Ganley, 2005) ; IGS2-R is synthesized toward the RFB site from the promoter near an origin of DNA replication (Houseley et al. 2007).

The IGS1-F transcripts that are synthesized from E-pro toward the 5S rRNA gene were reduced by ∼40% in the *spt4*Δ mutant (Fig. 3F, 3G), although this difference was not statistically significant. The level of IGS1-R transcript was detected below detection limit of Northern blotting, making it difficult to quantify signal intensities accurately. Transcription of IGS2-R, although it was not synthesized from E-pro, was also suppressed by 40% in the *spt4*Δ strain, compared to WT, but this difference was again not statistically significant (Fig. 3H, 3I).

As shown above, overexpression of *FOB1* in the *ADH1p-FOB1* strain induces smearing of chr XII band to the degree similar to that seen in the *sir2*Δ mutant (Fig. 2D–2F). A previous study shows that overexpression of *FOB1* causes weakened silencing of a marker gene inserted in the IGS1 and proposes that over-abundance of Fob1 causes inappropriate interactions of Fob1 with the RENT complex (Buck et al., 2016), which is composed of Sir2, Net1, and Cdc14, is recruited to IGS1 and enhances gene silencing (Huang and Moazed, 2003). Consistent with the previous finding, overexpression of *FOB1* led to de-repression of transcription from E-pro, as the *ADH1p-FOB1* strain showed ∼4-fold increase in the IGS1-F transcripts, compared to wild-type cells (Fig. 3J, 3K, WT/*FOB1* vs. WT/*ADH1p-FOB1*). As a cause of this phenotype, we identified that the Sir2 protein level was reduced by ∼85% in the *ADH1p-FOB1* cells, compared to the WT (*FOB1*) cells (Fig. 3L, 3M), indicating that Fob1 overexpression causes a reduction in the Sir2 protein level, enhancing transcription from E-pro. Deletion of *SPT4* from the *ADH1p-FOB1* strain did not alter the Sir2 protein level but caused a decrease in the level of the IGS1-F transcripts by ∼4-fold, compared to the *ADH1p-FOB1* strain (Fig. 3J–3M). Taken together, these findings indicate that Spt4 functions to enhance transcription from E-pro.

The IGS1-F and IGS2-R transcripts were increased by ∼7- and 8-fold in the *sir2*Δ mutant, respectively, compared to wild-type cells (Fig. 3F-3I). Thus, Sir2 regulates transcription of non-coding RNA not just from E-pro but also from the promoter in the IGS2. Deletion of *SPT4* from the *sir2*Δ mutant did not result in a decrease in transcription activity that was significant or comparable to that observed in the *SIR2* background (Fig. 3F–3I). The possibility that Spt4 represses *SIR2* expression would explain the similar phenotype for the *sir2*Δ *spt4*Δ double and *sir2* single mutant. However, this possibility was excluded, because the level of Sir2 was not altered by deletion of *SPT4* (Fig. 3L, 3M). Therefore, Spt4-mediated enhancement of transcription of non-coding RNA from E-pro depends on *SIR2*. Furthermore, the finding that only in *SIR2* cells the absence of Spt4 affected non-coding RNA transcription from E-pro is consistent with Spt4 dependence on *SIR2* for the regulation of rDNA stability.

### Spt4-mediated shortening of the replicative lifespan depends on *SIR2* and *FOB1*

The lifespan of the *spt4*Δ mutant is reported to be longer than that of the wild type, but the mechanism causing this remains unknown (Smith et al., 2008; McCormick *et al*., 2015). To explore how lifespan is extended in the *spt4*Δ mutant, we performed epistasis analysis of *SPT4* with known lifespan-modulating genes. Namely, we examined genetic interaction with *FOB1* that shortens lifespan and promotes rDNA instability and *SIR2* that extends lifespan and promotes rDNA stability (Defossez *et al*., 1999; Kaeberlein *et al*., 1999). Consistent with previous findings, the *sir2*Δ and *fob1*Δ mutants had shortened and extended lifespans, respectively (Fig. 4A, 4B). Deletion of *SPT4* led to an extension of lifespan in the wild-type background, consistent with previous findings (Smith *et al*., 2008; McCormick *et al*., 2015). This lifespan extension was comparable to that of cells lacking Fob1. However, deletion of *SPT4* from the *fob1*Δ background did not result in lifespan extension (Fig. 4A, 4B). Similarly, deletion of *SPT4* from the *sir2*Δ background did not lead to extension of lifespan (Fig. 4A, 4B). These findings indicate that Spt4 acts in the same pathway as Sir2 and Fob1 during the regulation of senescence.

**Figure 4.**
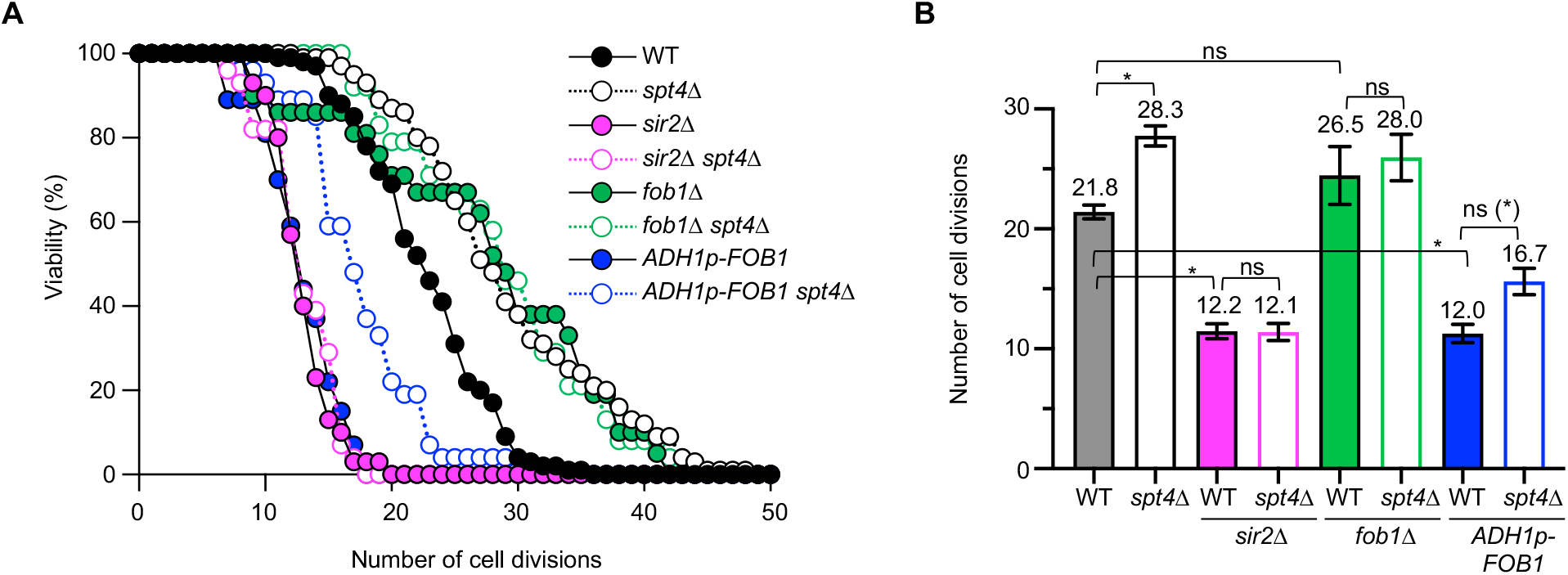
Spt4-mediated shortening of replicative lifespan depends on *FOB1* and *SIR2*. **(A, B)** Lifespans were determined by counting the number of daughter cells produced by a mother cell. Survival curve (A) and the mean lifespans (B) of the indicated strains were determined. The numbers of cells analyzed for lifespan measurements were n=101 (WT), 107 (*spt4*Δ), 33 (*sir2*Δ), 31 (*spt4*Δ *sir2*Δ), 24 (*fob1*Δ), 27 (*spt4*Δ *fob1*Δ), 26 (*ADH1p-FOB1*), and 26 (*ADH1p-FOB1 spt4*Δ). Statistical significance was determined by a Kruskal-Wallis non-parametric test with Dunn’s multiple comparisons tests. An asterisk indicates a significant difference (*p* < 0.001); ns marks a difference that is not statistically significant. ns (*) between *ADH1p-FOB1* and *ADH1p-FOB1 spt4*Δ strains indicates that the difference in the mean lifespan between *ADH1p-FOB1* and *ADH1p-FOB1 spt4*Δ strains is not statistically significant by a Kruskal-Wallis non-parametric test with Dunn’s multiple comparisons tests but was found to be significant by the Mann-Whitney, nonparametric test (*P* < 0.001).

### Spt4 restricts replicative lifespan through stimulation of non-coding RNA transcription from E-pro

E-pro activity induces rDNA instability and cellular senescence (Kobayashi and Ganley, 2005; Saka et al., 2013). Indeed, enhanced transcription of non-coding RNA in the *ADH1p-FOB1* strain was accompanied with shortening of replicative lifespan, compared to the wild-type strain (Fig. 3J, 3K and Fig. 4). Furthermore, deletion of *SPT4* from the *ADH1p-FOB1* strain led to reduced transcription of non-coding RNA and rDNA instability (Fig. 2D–2F, 3J, 3K). The *ADH1p-FOB1 spt4*Δ strain clearly showed extended lifespan on the survival curve, compared to the *ADH1-FOB1* strain (Fig. 4A). It should be noted that although the multiple comparisons tests did not show statistical significance in the mean lifespan between *ADH1p-FOB1* and *ADH1p-FOB1 spt4*Δ strains (Fig. 4B), the difference was statistically significant (*P* < 0.001 in the Mann-Whitney, nonparametric test). These results point to the possibility that reduced transcription from E-pro may be a primary determinant of lifespan extension in the absence of Spt4. To test this possibility, we used the Gal-pro strain (Fig. 5A), in which the E-pro is replaced by the galactose-inducible bi-directional *GAL1/10* promoter in all rDNA copies (Kobayashi and Ganley 2005). In the Gal-pro strain, transcription of noncoding RNA is repressed in glucose-containing media but it is induced in medium with galactose as the carbon source (Kobayashi and Ganley, 2005) (Fig. 5B). In the Gal-pro strain, the rDNA is stabilized and lifespan is extended when cells are grown in the presence of glucose, compared to cells cultured in galactose-containing media (Kobayashi and Ganley, 2005; Saka *et al*., 2013).

**Figure 5.**
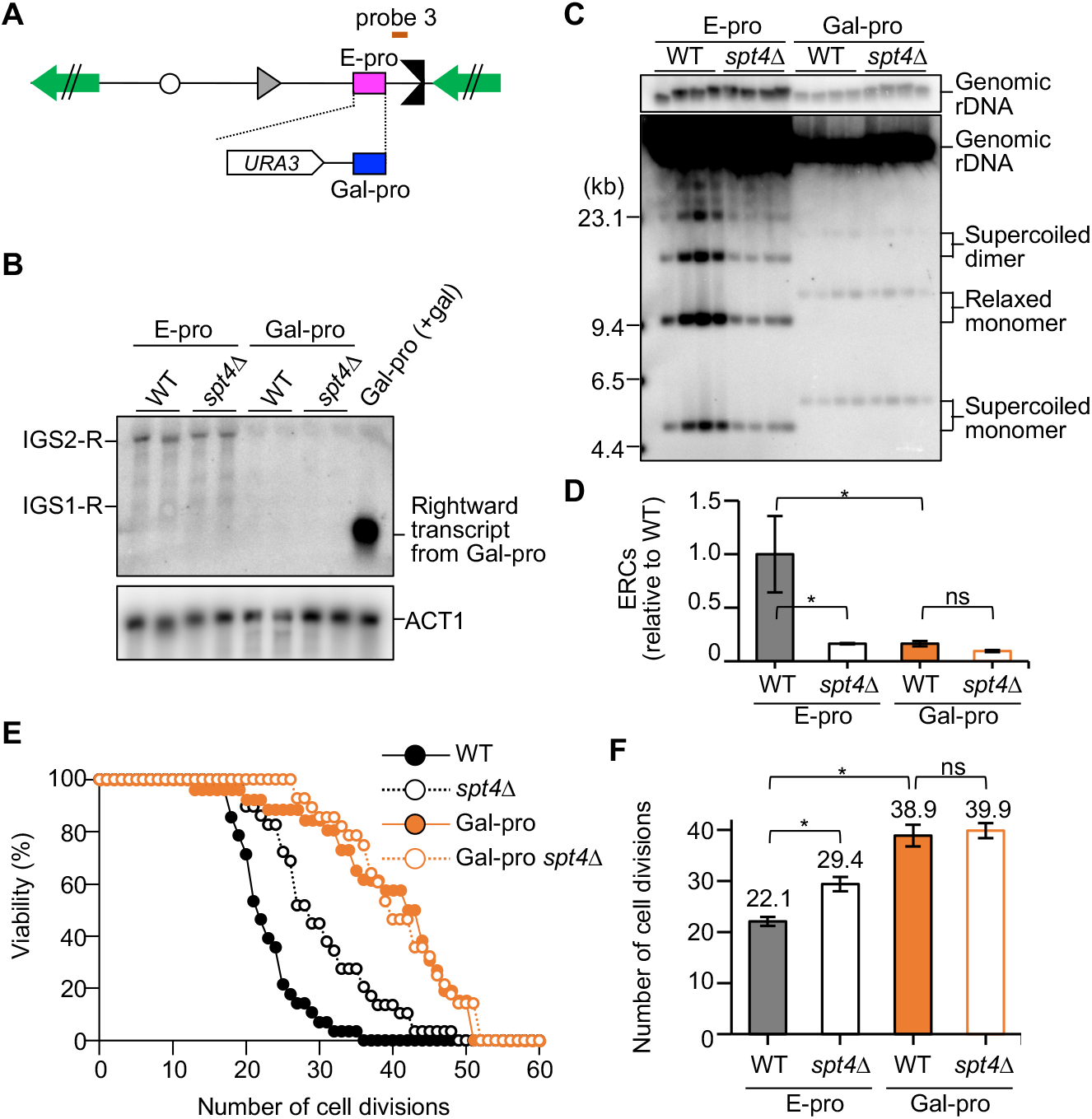
Spt4-mediated rDNA instability and shortening of replicative lifespan depends on transcription activity from E-pro. **(A)** Replacement of the E-pro with the bi-directional *GAL1/10* promoter in all the rDNA copies in the Gal-pro strain. Probes 3 indicates the position of the single-stranded probe used for detection of non-coding RNA in (B). **(B)** Detection of non-coding RNAs transcribed from E-pro and Gal-pro. The indicated strains were cultured in media with glucose, except for control cells that were grown in the presence of galactose (+gal). RNA was isolated and separated in formaldehyde-containing agarose, followed by Northern blotting. Transcripts synthesized toward the RFB site were detected with probe 3 (see (A)). Membranes were reprobed with a probe hybridizing to *ACT1* transcripts (ACT1). **(C)** ERC detection. DNA isolated from the indicated strains grown in glucose-containing media was separated by agarose gel electrophoresis, followed by Southern blotting with probe 1 (see Figs. 1A, 3A). Genomic rDNA and supercoiled and dimer forms of monomeric and dimeric ERCs are indicated. Sizes of lambda DNA-Hind III markers are indicated. **(D)** Quantitation of ERC levels. Levels of ERCs in (C) were determined by the ratio of the sum of monomeric and dimeric ERCs relative to genomic rDNA, which was normalized to the average of wild-type clones (bars show mean ± s.e.m). One-way ANOVA was performed for multiple comparisons (asterisks indicate statistically significant difference between wild type and mutants [*p* < 0.05]; ns indicates no significant difference (*p* > 0.05)). **(E, F)** Lifespans were determined by counting the number of daughter cells produced by mother cells growing on medium with glucose. Survival curve (E) and the mean life spans (F) of the indicated strains are shown. The numbers of cells analyzed for lifespan measurements were n=28 (E-pro), 29 (E-pro *spt4*Δ), 26 (Gal-pro), and 28 (Gal-pro *spt4*Δ). Statistical significance was determined by a Kruskal-Wallis non-parametric test with Dunn’s multiple comparisons tests. An asterisk indicates a significant difference (*p* < 0.05); ns marks a difference that is not statistically significant (*p* > 0.05).

In the presence of glucose, non-coding RNA transcription was kept at a level below the detection limit of Northern blotting in both Gal-pro and Gal-pro *spt4*Δ strains (Fig. 5B). Furthermore, the level of ERCs was extremely low in the Gal-pro strain, compared to the E-pro strain, and this level did not alter in the absence of Spt4 (Figs. 5C, 5D, lanes Gal-pro *spt4*Δ). The lifespan of the Gal-pro strain was longer than that of the E-pro strain, as described previously (Saka et al. 2013). Although deletion of *SPT4* led to an extension of the replicative lifespan of E-pro cells, this deletion did not further extend the replicative lifespan of Gal-pro cells (Fig. 5E, 5F). Therefore, transcription of non-coding RNA depending on the presence of Spt4 induces shortening of the replicative lifespan, which is connected to rDNA stability.

### Spt4 protein level increases in old cells

A previous study demonstrated that the Sir2 protein level decreases by ∼80% in cells that have divided ∼6 times, which is in line with its anti-aging function (Fine *et al*., 2019). We also examined the alteration in protein levels by comparing ’old’ cells grown for 7 to 8 divisions, to ’young’ cells that had not divided or just once. To this end, we tagged the ORFs of *SIR2* or *SPT4* with a triple HA epitope and separated young from old cells using magnetic streptavidin beads that only bound to cell-surface proteins of old cells after their initial exposure to a biotinylation reagent. Tagged protein levels were assessed by Western blotting. Using this approach, we also detected a statistically significant reduction in Sir2 levels as cells age (Fig. 6A, 6B). Possibly due to experimental differences, the ∼40% reduction we observed was less than reported.

**Figure 6.**
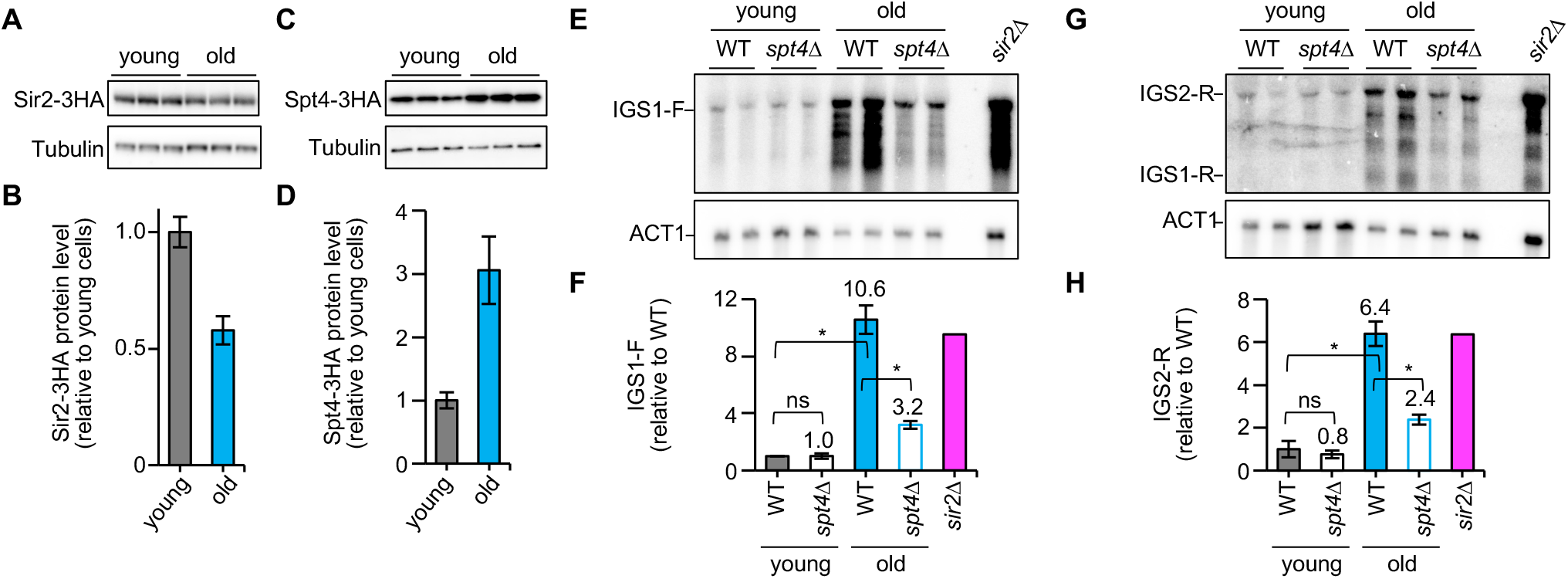
The function of Spt4 becomes more important as cells age. **(A, C)** Detection of Sir2 and Spt4 protein levels. Strains were constructed expressing Sir2 or Spt4 protein with a triple HA epitope at the C-terminus. Protein extracts isolated from three independent clones of the indicated strains were separated by SDS-PAGE, followed by Western blotting with antibodies against HA to detect Sir2-3HA (A) or Spt4-3HA (C), and against tubulin. **(B, D)** Quantitation of Sir2 and Spt4 expression. The level of Sir2-3HA (B) and Spt4-3HA (D) protein was quantified relative to tubulin, which was normalized to the average of their levels in young cells (bars show mean ± s.e.m). **(E, G)** Synthesis of non-coding RNA in young and old cells. In each of the three independent cultures, young cells were split from old cells for the indicated strains. The control *sir2*Δ cells were collected from one exponentially growing culture. RNA was isolated and separated by gel electrophoresis over formaldehyde-containing agarose, followed by Northern blotting. Transcripts synthesized toward the 5S (IGS1-F) and RFB sides (IGS2-R) were detected with probes 4 and 3 (see Fig. 3A), respectively. Membranes were reprobed with the *ACT1* probe. RNA from the *sir2*Δ strain was isolated from unsorted young cells. **(F, H)** Quantitation of non-coding RNA levels. The signals of non-coding RNA transcripts (E, G) were quantified relative to *ACT1*, which was normalized to the average of wild-type clones (bars show mean ± s.e.m). One-way ANOVA was performed for multiple comparisons for samples, except for *sir2*Δ (asterisks indicate statistically significant difference between wild type and mutants [*p* < 0.05]; ns indicates no significant difference (*p* > 0.05)).

If Spt4 is an aging factor, it is feasible that its protein level increases as cells undergo more cell divisions. In agreement with this hypothesis, we found that the level of HA-tagged Spt4 protein was elevated in old cells by ∼3-fold, when this was compared to the abundance of Spt4 in young cells (Fig. 6C, 6D). As reported previously (Pal et al., 2018), in old cells we found that the levels of transcripts synthesized from E-pro increased by >6-fold in either direction (Fig. 6E–6H). Although transcript levels did not change in young cells after deletion of *SPT4*, they were >3-fold lower in old *spt4*Δ cells, compared to old wild-type cells (Fig. 6F, 6H). Thus, the function of Spt4 appears to become more important as cells age. We conclude that Spt4 is a strong candidate for an aging factor that restricts the lifespan of cells, in this case by elevated activation of non-coding RNA transcription as cells undergo more cell-divisions. This drives rDNA instability and senescence gets triggered.

## Discussion

In the budding yeast, the rDNA locus is thought to be a source of a putative aging signal. Thus, the maintenance of rDNA stability is quite important for enhanced longevity. We identified that Spt4 is a key factor that accelerates rDNA instability and cellular senescence by enhancing transcription of non-coding RNA from E-pro (Fig. 1–4). The amount of Spt4 was increased in aging cells and its depletion extended the lifespan (Fig. 4, 6). By genetic analysis, Spt4 was demonstrated to function downstream of Fob1, Sir2, and E-pro activity in the regulation of rDNA stability and replicative lifespan (Fig. 2, 4, 5). These findings suggest that Spt4 works as an aging factor in budding yeast.

Our analysis of the *spt4*Δ mutant revealed that the activity of Spt4 stimulates transcription of non-coding RNA from E-pro and an upstream promoter in the IGS2, destabilizing the rDNA and extending replicative lifespan (Fig. 1–4). These Spt4*-*dependent phenotypes completely disappeared when cells lacked Sir2 and Fob1 (Fig. 2–4). When transcription from E-pro is repressed in the Gal-pro strain, Spt4 activity had no effect on rDNA stability or lifespan in this strain (Fig. 5). These results strongly suggest that the function of Spt4 to activate transcription from E-pro is a direct cause of accelerating cellular senescence through rDNA instability.

Based on our findings and in view of the enzymatic activities of Sir2 and Spt4, we developed a working hypothesis how Spt4 enhances E-pro transcription in a Sir2 dependent manner (Fig. 7). Sir2 deacetylates the histones around E-pro in the wild-type strain (“+Sir2”) which results in compacti jon of the chromatin (Fritze et al., 1997; Huang and Moazed, 2003). The nucleosome can become a barrier to progression of RNAP II (Ehara and Sekine, 2018; Kujirai and Kurumizaka, 2020). Thus, closed chromatin structures formed by Sir2 causes pausing of the RNAP II transcription machinery. The activity of the elongation factor Spt4-Spt5 reduces this pause or releases this paused RNAP II, increasing the elongation rate of RNAP II at the usual rate (“+Spt4”). Production of non-coding RNA triggers rDNA instability (possibly by displacement of cohesin), which restricts the lifespan. In the absence of Spt4 (“-Spt4”), transcription elongation activity is reduced, resulting in a decrease of non-coding RNA synthesis that contributes to rDNA stability and a lifespan extension. In a *SIR2* defective background (“- Sir2”), the histones remain acetylated and transcription by RNAP II can proceed normally regardless of Spt4 activity (“±Spt4”) with elevated rDNA instability and a shortened lifespan as the outcome.

**Figure 7.**
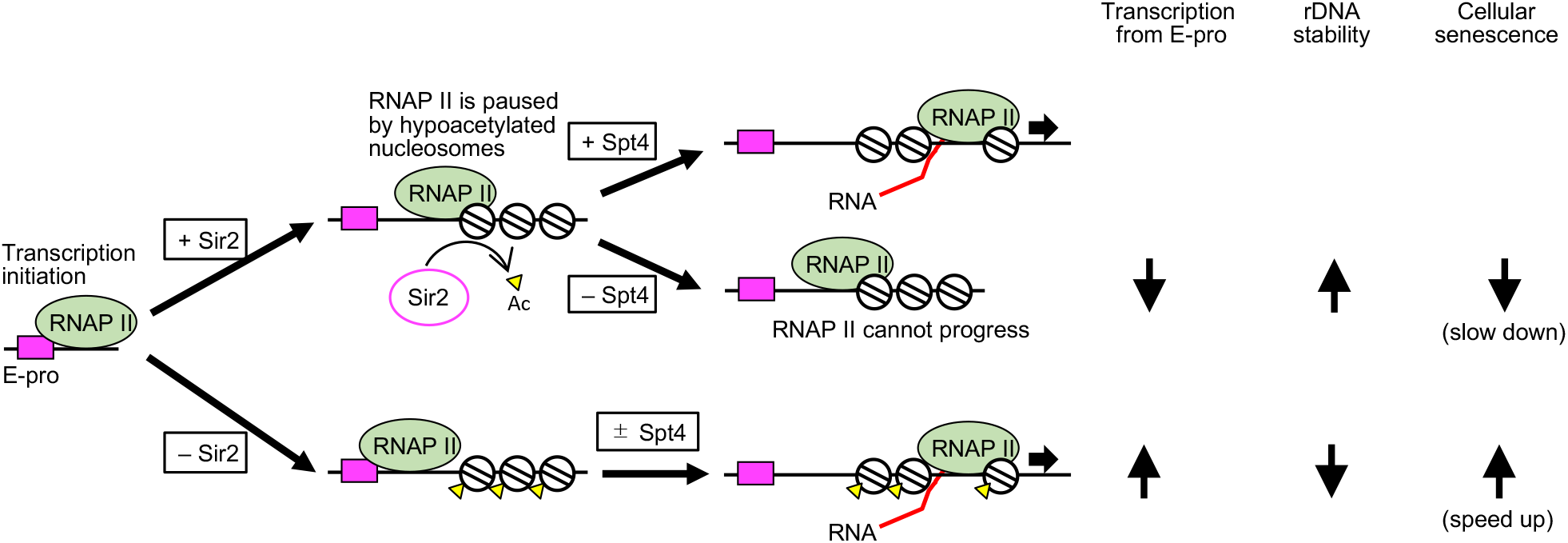
Spt4 promotes transcription of non-coding RNA from E-pro in a manner dependent on Sir2. In wild-type cells (+Sir2, +Spt4), due to the histone deacetylase activity of Sir2, histones lose their acetyl modifications (Ac) and closed chromatin structures are formed around E-pro. Such chromatin structures block progression of RNA polymerase II (RNAP II). The Spt4-Spt5 complex is a transcription elongation factor that facilitates progression of RNAP II, leading to transcription of non-coding RNA at a normal rate when histones are deacetylated. Thus, in cells carrying Sir2 but not Spt4 (or defective Spt5) (+Sir2, –Spt4), RNAP II cannot progress normally and non-coding RNAs are transcribed at a reduced rate, which leads to an increase in rDNA stability, generating less aging signal and slows down cellular senescence. In the absence of Sir2, chromatin structures are kept open, stimulating transcription of non-coding RNA by RNAP II, regardless of the presence or absence of Spt4-Spt5 activity (+Sir2, ±Spt4). The increased transcription activity triggers rDNA instability, accelerating cellular senescence. As cells age, the protein level of Spt4 increases, elevating transcription from E-pro, rDNA instability and cellular senescence. In old cells lacking Spt4, the elongation activity is reduced and the transcription is decreased from E-pro (-Spt4), leading to rDNA stability and extension of lifespan.

Sir2 is classified as an anti-aging factor, because its absence causes lifespan shortening, its over-production leads to extension of lifespan, and its protein level is reduced during normal aging (Kaeberlein *et al*., 1999; Fine *et al*., 2019). The activity of Spt4 seems to have an opposite effect on aging. Deletion of *SPT4* extended the lifespan by ∼30% (Figs. 4, 5E, 5F), as previously reported (Smith *et al*., 2008; McCormick *et al*., 2015). Furthermore, we observed that as cells age, expression of Spt4 increased over 3-fold, which was accompanied with the significantly elevated levels of E-pro-derived transcripts of over 10-fold when cells age (Fig. 6). An increase in Spt4 levels, together with reduced Sir2 levels, may be instrumental in enhancing E-pro transcription by stimulating RNAP II processivity. However, a change of Sir2 distribution with an impact on lifespan has previously been described: Sir2 relocalizes to telomeres and the silent mating-type loci in the absence of Rif1, a mutation that affects lifespan (Salvi et al., 2013). Therefore, it is also feasible that although Spt4 does not influence the Sir2 protein level (Fig. 3L, 3M), it may be involved in Sir2 distribution in the genome. An increased abundance of Spt4 in old cells may prevent localization of Sir2 to the IGS1. Then, non-coding RNA is transcribed from E-pro in more rDNA copies as cells age, compared to the situation in young cells, leading to enhanced transcript levels in old cells.

So far, experimental evidence is lacking that would explain how Spt4 protein level increases in aging cells. A possible mechanism is that Sir2 represses *SPT4* expression. If this is the case, Spt4 works downstream of Sir2 which fits the results of our genetic analysis on lifespan (Fig. 4), DSB formation and the regulation of expression and elongation of non-coding transcripts (Fig. 3), where Spt4 activity depended on that of Sir2. Whether 40% reduction in the levels of Sir2 we detected in aging cells would be sufficient to accomplish this, will have to be tested more specifically.

How do Spt4-dependent transcription activation in the rDNA IGSs induce cellular senescence? First, previous studies propose that the accumulation of ERCs induces cellular senescence by titrating factors that are important for transcription, genome maintenance, and cell growth (Sinclair and Guarente, 1997; Kwan et al., 2013; Neurohr et al., 2018). In line with this model, Spt4-mediated E-pro activation induces ERC formation, potentially driving cellular senescence (Fig. 1B–1D, 2E, 2F, 5C, 5D). The second model centers around the rDNA stability induced by DNA damage in the rDNA (Ganley *et al*., 2009; Kobayashi, 2011; Ganley and Kobayashi, 2014). The Spt4 activity promotes DSB formation at arrested forks, although its impact on DSB end resection, an initiating event of rDNA copy number changes, could not be assessed in our study (Fig. 3D, 3E). The Spt4-mediated transcription activation of E-pro may promote homologous recombination-mediated rDNA instability by promoting DSB formation and its end resection, which needs to be clarified in future studies. Lastly, non-coding RNA transcription may drive cellular senescence in as yet unidentified manners. The lifespan of the Gal-pro strain in glucose was markedly longer than that of the wild-type strain (38.9 to 22.1 divisions, Figs. 5E, F), and even longer than that of the *fob1* mutant (26.5) that does not generate DSBs at the RFB site (Fig. 4). Furthermore, a noticeable upregulation of pre-rRNA synthesis had been observed as a concomitant of ERC accumulation preceding senescence (Morlot et al., 2019). Therefore, these observations suggest that noncoding transcription itself could be part of a pathway leading to senescence. The above models relating rDNA maintenance to aging do not need to be mutually exclusive, as might be revealed by further research into the molecular mechanisms controlling the events marked by DSB formation, the accumulation of ERCs and non-coding RNA and how these contribute to senescence.

Compared to wild type, the replicative lifespan of Gal-pro strains (including the Gal-pro *spt4*Δ mutant) was extended by ∼ 80%, while the lifespan of the *spt4*Δ mutant with the native E-pro was increased by ∼30% (Figs. 4, 5E, 5F). These differences indicate that transcription from E-pro is not the only factor that determines the onset of senescence via the rDNA. Other candidates than Spt4, which we count as an E-pro-mediated aging factor, have been identified (Fig. 1B). How the increase in rDNA stability observed in the absence of these putative aging factors, extends replicative lifespan will be analyzed in our future studies. The proteins encoded by *SIR2* and *SPT4* are conserved in mammalian cells. Therefore, the findings we reported here will be relevant for understanding the molecular mechanisms that link genomic instability in the rDNA to the initiation of senescence signals in human cells.

### AUTHOR CONTRIBUTIONS

M.Y., M.S. and T.K. designed the experiments, analyzed the data and wrote the paper. M.Y. performed the experiments.

## ACKNOWLEDGEMENTS

We are grateful to David Schneider for yeast strains. This work was supported by JST CREST (grant number JPMJCR19S3 to T.K.), JST FOREST Program (grant number JPMJFR214P to M.S.), and Grant-in-Aid for Scientific Research (17H01443 and 21H04761 to T.K, and 20H05382 to M.S.).

## MATERIALS AND METHODS

### Yeast Strains and Culture Methods

Yeast strains used in this study are derivatives of W303 (*MATa ade2-1 ura3-1 his3-11, 15 trp1-1 leu2-3,112 can1-100*). The *spt5-C292R* and *spt5-S324P, E427K* strains used in Fig. 1D were previously constructed and a kind gift from David Schneider (Anderson *et al*., 2011). Gene tagging or deletion was performed by standard one-step gene replacement methods and the mutant strains were confirmed by PCR-based genotyping. Prior to use, strains were streaked from a glycerol stock onto YPD plates (1% w/v yeast extract, 2% w/v peptone, 2% w/v glucose, and 2% w/v agar). Yeast cells were grown at 30°C, except for the experiment in Fig. 1C where cells were grown at 27°C. For each DNA, RNA or protein preparation, 5 ml of YPD was inoculated with a single colony and incubated overnight. For PFGE and ERC analyses, overnight cultures were collected (5 × 10^7^ cells/plug) and washed twice with 50 mM EDTA pH 7.5. For two-dimensional (2D) agarose gel electrophoresis and DSB analyses, overnight cultures were diluted into 50 mL of YPD medium to OD_600_ = 0.1 and grown until OD_600_ = 0.4. The cultures were immediately treated with 1/1,000 vol of 10% sodium azide, collected (5 × 10^7^ cells/plug) by centrifugation for 2 min at 2,400 × *g* at 4°C and washed twice with 50 mM EDTA pH 7.5. To prepare RNA, overnight cultures were diluted into 15 mL of YPDA medium (YPD containing 40 μg/mL adenine sulfate) to an OD_600_ = 0.1and collected when the cultures reached an OD_600_ of 0.8. Protein extracts were prepared from 3 × 10^7^ cells grown in 50 mL of YPD medium to OD_600_ = 0.4 after dilution of overnight cultures to OD_600_ = 0.1.

### Genomic DNA Preparation

For PFGE, ERC, DSB, and 2D gel analyses, genomic DNA was prepared in low melting temperature agarose plugs as described previously (Sasaki and Kobayashi, 2017; 2021). Briefly, collected cells were resuspended in 50 mM EDTA pH 7.5 (33 μL cell per 5 × 10^7^ cells and incubated at 42°C. For each plug, 33 μL cell suspension was mixed with 66 μL solution 1 (0.83% low-melting-point agarose SeaPlaque GTG (Lonza), 170 mM sorbitol, 17 mM sodium citrate, 10 mM EDTA pH 7.5, 0.85% β-mercaptoethanol, and 0.17 mg/mL Zymolyase 100 T (Nacalai)), poured into a plug mold (Bio-RAD) and placed at 4°C for the agarose to be solidified. Plugs were transferred to a 2 mL-tube containing solution 2 (450 mM EDTA pH 7.5, 10 mM Tris-HCl pH 7.5, 7.5% β-mercaptoethanol and 10 μg/mL RNaseA (Macherey-Nagel)), and incubated for 1 h at 37°C. Plugs were then incubated overnight at 50°C in solution 3 (250 mM EDTA pH 7.5, 10 mM Tris-HCl pH 7.5, 1% sodium dodecyl sulfate (SDS) and 1 mg/mL Proteinase K (Nacalai)). Plugs were washed four times with 50 mM EDTA pH 7.5 and stored at 4°C in 50 mM EDTA pH 7.5.

### Southern blotting

#### Agarose gel electrophoresis

##### ERC assay

ERC assay was performed as described previously (Goto et al., 2021; Sasaki and Kobayashi, 2021). Half an agarose plug was placed on a tooth of the comb. After the comb was set in the gel tray (15 × 25 cm), 300 ml of 0.4% agarose (SeaKem LE Agarose, Lonza) in 1x TAE (40 mM Tris base, 20 mM acetic acid, and 1 mM EDTA pH 8.0) was poured into the tray and allowed to set, after which 500 ng of lambda *Hind* III DNA marker was applied to an empty lane. The electrophoresis was run on a Sub-cell GT electrophoresis system (Bio-Rad) in 1.5 L of 1x TAE at 1.0 V/cm for ∼48 h at 4°C with buffer circulation. The buffer was changed every ∼24 h. DNA was stained with 0.5 μg/mL EtBr for 30 min and then photographed.

##### 2D gel electrophoresis

2D gel electrophoresis was performed as described previously with slight modifications (Goto *et al*., 2021). One-half of an agarose plug was placed in a 2-mL tube. The plug was equilibrated twice in 1 mL of 1x M buffer (TaKaRa) by rotating the tube for 30 min at room temperature. After discarding the buffer, the plug was incubated in 160 μL of 1x M buffer containing 160 units of *Nhe* I (TaKaRa) overnight at 37°C. The plug was placed on a tooth of the comb. The comb was set in the gel tray (15 × 25 cm), 0.4 % agarose solution in 1x TBE was poured and the gel was allowed to solidify. 600 ng of lambda *Hind* III DNA markers were applied to the empty lane. The electrophoresis was performed on a Sub-cell GT electrophoresis system (Bio-Rad) in 1.5 L of 1x TBE at 1.32 V/cm for 14 h at room temperature with buffer circulation.

After electrophoresis completed, DNA was stained with 0.3 μg/mL EtBr for 30 min and then photographed. Gel slices containing DNA ranging from 4.7 to 9.4 kb were excised, rotated 90°, and placed onto the gel tray used for the second agarose gel electrophoresis for which 1.2 % agarose solution in 1x TBE containing 0.3 μg/mL of EtBr was poured into the tray. The second-dimension electrophoresis was run on a Sub-cell GT electrophoresis system (Bio-Rad) in 1.5 L of 1x TBE containing 0.3 μg/mL of EtBr at 6.0 V/cm for 6 h at 4°C with buffer circulation. After electrophoresis, the DNA was photographed.

##### DSB assay

The DSB assay was performed as described previously (Sasaki and Kobayashi, 2017; 2021). One-third of an agarose plug was cut and placed in a 2-mL tube. The plug was equilibrated four times in 1 mL of 1x TE (10 mM Tris base pH 7.5 and 1 mM EDTA pH 8.0) by rotating the tube for 15 min at room temperature. The plug was equilibrated twice in 1 mL of 1x NEBuffer 3.1 (New England Biolabs) by rotating the tube for 30 min at room temperature. After discarding the buffer, the plug was incubated in 160 μL of 1× NEBuffer 3.1 buffer containing 160 units of *Bgl* II (New England Biolabs) overnight at 37°C. The plug was placed on a tooth of the comb which was set into the gel tray (15 × 25 cm), into which 0.7 % agarose solution in 1x TBE was poured; after setting of the gel 600 ng of lambda *Bst* ELJ DNA markers were applied to an empty lane. The electrophoresis was run on a Sub-cell GT electrophoresis system (Bio-Rad) in 1.5 L of 1x TBE at 2.0 V/cm for 22 h at room temperature with buffer circulation. After electrophoresis, the DNA was stained with 0.5 μg/mL EtBr for 30 min and then photographed.

#### DNA transfer

After agarose gel electrophoresis, the gel was incubated by gentle mixing in 500 mL of 0.25 N HCl for 30 min, 500 mL of denaturation solution (0.5 N NaOH, 1.5 M NaCl) for 30 min, and in 500 mL of transfer buffer (0.25 N NaOH, 1.5 M NaCl) for 30 min. DNA was transferred to Hybond-XL (GE Healthcare) by the standard capillary transfer method with transfer buffer. DNA was fixed to the membrane by soaking the membrane in 300 mL freshly prepared 0.4 N NaOH for 10 min with gentle shaking, followed by rinsing the membrane with 2x SSC for 10 min. The membrane was dried and stored at 4°C.

#### Probe preparation

Probes were prepared as described previously (Sasaki and Kobayashi, 2017; 2021). Double-stranded DNA fragments were amplified by PCR. Probe 1 used for ERC and 2D analyses was amplified with the primers 5’-CATTTCCTATAGTTAACAGGACATGCC and 5’- AATTCGCACTATCCAGCTGCACTC; and probe 2 used for DSB analysis was amplified with the primers 5’-ACGAACGACAAGCCTACTCG and 5’-AAAAGGTGCGGAAATGGCTG. A portion of PCR products was gel-purified and seeded for a second round of PCR with the same primers. PCR products were gel-purified and 50 ng was used for random priming reactions in the presence of radio-labeled nucleotide, [α-^32^P]-dCTP (3,000 Ci/mmol, 10 mCi/ml, Perkin Elmer), using Random Primer DNA Labeling Kit (TaKaRa), according to the manufacturer’s instructions. Unincorporated nucleotides were removed using ProbeQuant G-50 Micro Columns (GE Healthcare). The radio-labeled probes were heat-denatured for 5 min at 100°C, immediately prior to hybridization to the membrane.

#### Hybridization

Southern hybridization was performed as described previously (Sasaki and Kobayashi, 2017; 2021). The membrane was pre-wetted with 0.5 M phosphate buffer pH 7.2 and pre-hybridized for 1 h at 65°C with 25 mL of hybridization buffer (1% bovine serum albumin, 0.5 M phosphate buffer pH 7.2, 7% SDS, 1 mM EDTA pH 8.0). After buffer was discarded, the membrane was hybridized with 25 mL of hybridization buffer and heat-denatured probe overnight at 65°C. The membrane was washed four times for 15 min at 65°C with wash buffer (40 mM phosphate buffer pH 7.2, 1 % SDS, 1 mM EDTA pH 8.0) and exposed to a phosphor screen.

#### Image analysis

Membranes were exposed to a phosphor screen for several days to achieve a high signal to noise ratio for DNA molecules that are low in abundance such as ERCs, replication intermediates in 2D assays, and arrested forks and DSBs. The radioactive signal was detected using a Typhoon FLA 7000 (GE Healthcare). In ERC assays, genomic rDNA signal was used for normalization of ERC signals. Thus, the membranes were re-exposed to the phosphor screen for a short time before this signal reached saturation. ERC bands and genomic rDNA were quantified using FUJIFILM Multi Gauge version 2.0 software (Fujifilm) and scans from long and short exposures, respectively. The levels of ERCs were calculated by dividing the sum of signal intensities of ERCs with those of genomic rDNA. When the levels of ERCs were compared between samples loaded on different gels, pieces of the plug excised from a specific DNA sample were loaded on different gels, blotted and hybridized. Signal of these plugs was used for normalization of samples on different gels. In 2D analyses, bubbles, Y arcs, RFB spots, and double Y spots were quantified using ImageJ (NIH). The RFB activity was calculated as the ratio between the signal for the spot with arrested forks and the sum of signals encompassing the total of DNA replication intermediates. Signals of DSBs and arrested forks were quantified using FUJIFILM Multi Gauge version 2.0 software (Fujifilm). The DSB frequency was calculated by normalizing the DSB signal to that of the arrested forks.

### PFGE

PFGE was performed as described previously (Sasaki and Kobayashi, 2017; 2021) . Briefly, one-third of a plug was placed on a tooth of the comb, including a piece with *Hansenula wingei* chromosomal DNA markers (Bio-Rad). The comb was set into the gel tray, and 1.0% agarose solution was poured (Pulsed Field Certified Agarose, Bio-Rad) in 0.5x TBE (44.5 mM Tris base, 44.5 mM boric acid and 1 mM EDTA pH 8.0). PFGE was run on a Bio-Rad CHEF DR-III system in 2.2 L of 0.5x TBE under the following conditions: 3.0 V/cm for 68 h at 14°C, 120° included angle, initial switch time of 300 s, and final switch time of 900 s. After electrophoresis, DNA was stained with 0.5 μg/mL ethidium bromide (EtBr) for 30 min, washed with dH_2_O for 30 min, and then photographed.

### Preparation of protein extracts

Protein extracts were prepared from yeast cells, as described previously with a slight modification (Iida and Kobayashi, 2019). Cells were re-suspended in 500 μL of ice-cold dH_2_O and 75 μL of Alkali-2ME solution (1.295N NaOH and 7.5% v/v 2-mercaptoethanol) by vortexing and incubated on ice for 10 min. Cell suspensions were mixed with 75 μL of 50% w/v Trichloroacetic Acid by vortexing and incubated on ice for 10 min. After centrifugation at 10,000 × *g* for 5 min at 4°C, the supernatant was completely removed and the pellets were resuspended in 50 μL of buffer (800 μl of 200 mM Tris-HCl pH 8.8 and 200μl of 5x SDS-loading buffer (250 mM Tris-HCl, pH 6.8, 8% SDS, 0.1% bromophenol blue, 40% glycerol, 100 mM Dithiothreitol)). Protein concentration was quantified using a Bradford assay (Bio-Rad). Protein samples were stored at -20°C.

#### *Western* blotting

Protein samples were prepared by mixing 20–25 μg of total protein extracts with 1x SDS-loading buffer to an end volume of 10 μL. Protein samples were boiled for 5 min at 65°C, and centrifugated at 20,000 × *g* for 5 min at 4°C. Supernatants were loaded and separated on 7.5% or 5–20% pre-made polyacrylamide gel (Atto) in 400 mL of 1x SDS-PAGE running buffer (25 mM Tris, 192 mM Glycine, 0.1% SDS). Proteins were transferred to Immobilon-P PVDF membrane (Merck-Millipore) in 1x transfer buffer (25 mM Tris, 192 mM glycine, 10% methanol) for 60 min at 100 V at 4°C using a Mini Trans-Blot cell (Bio-Rad). After transfer, the membrane was blocked in 5% Skim Milk (Wako) in PBS-T (1x PBS, 0.05% Tween20) at room temperature for 1 h. The membrane was incubated with HRP-conjugated antibodies overnight at 4°C: anti-3HA-HRP (F-7) (Santa Cruz [sc-7392], 1:5,000 in blocking buffer with 1% skim milk) and anti-tubulin-HRP (YL1/2) (BIO-RAD [MCA77P], 1/5000 dilution in blocking buffer with 5% Skim Milk Powder). After the membranes were washed three times for 5 min with PBS-T, the membranes were incubated with the chemiluminescent substrate, Immobilon Western Chemiluminescent HRP Substrate (Merck-Millipore, WBKLS0100), according to the manufacturer’s instruction. Protein signals were detected with a Fusion imaging system (Vilber Lourmat). Protein signals were quantified using ImageJ (NIH).

### Yeast RNA preparation

RNA was prepared as described previously (Iida and Kobayashi, 2019), with a slight modification. Collected cells were resuspended in 400 μL of TES (10 mM Tris-HCl pH 7.5, 10 mM EDTA pH 7.5, 0.5% SDS) and 400 μl of acidic phenol by vortexing for about 10 s. Cells were incubated at 65°C for 1 h with occasional vortexing every 15 min. Cell suspensions were incubated for 5 min on ice, and centrifuged at 20,000 × *g* for 10 min at 4°C. The aqueous phase was transferred to a new tube, and mixed with an equal volume of acidic phenol by vortexing for 10 s. Samples were incubated for 5 min on ice and centrifuged at 20,000 × *g* for 10 min at 4°C. The aqueous phase was transferred to a new tube. Then, 1/10 vol. of 3M sodium acetate (pH 5.3) and 2.5 vol. of 100% ethanol were added and RNA was precipitated overnight at -20°C. After centrifugation at 20,000 × *g* for 10 min at 4°C, RNA was washed with 70% ethanol. RNA pellets were resuspended in 30 μL of dH_2_O treated with 0.1% diethylpyrocarbonate (DEPC). The concentration of RNA was quantified using a NanoDrop ND-1000 spectrophotometer (Thermo Fisher Scientific). RNA samples were stored at -80°C.

### Northern blotting

18∼30 μg of total RNA was brought up to 7 μL with DEPC-treated dH_2_O and mixed with 17 μL of RNA sample buffer (396 μL deionized formamide, 120 μL 10 x MOPS buffer [0.2 M MOPS, 50 mM sodium acetate pH 5.2, 10 mM EDTA pH 7.5 in DEPC-treated dH_2_O], 162 μL 37% formaldehyde). Samples were heated at 65°C for 20 min, followed by a rapid chill on ice. 6 μL of 6x Gel Loading Dye (B7025S, New England Biolabs) and 1.5 μL of 2.5 mg/ml EtBr were added to each sample. The 1% agarose gel was made by dissolving 1 g of agarose powder in 73 mL DEPC-treated dH_2_O and, after cooling to 60°C, addition of 17 mL 37% formaldehyde and 10 mL 10x MOPS buffer. The solution was poured into a gel tray (13 × 12 cm) and allowed to set; 10 μg of total RNA (10 μL) and 1.8 μg of DynaMarker RNA High markers were applied. Agarose gel electrophoresis on a Mupid EX system (Takara) was in 400 mL of 1x MOPS buffer at 20V for 20 min and then at 100V until the bromophenol blue dye migrated about 2/3 of the gel. After electrophoresis, the gel was photographed and RNA was transferred to Hybond-N+ (GE Healthcare) by standard capillary transfer.

Strand-specific probes were prepared from double-stranded DNA fragments, amplified by PCR and gel-purified. PCR primers for probe 3 (against IGS1-F) were 5’- AGGGAAATGGAGGGAAGAGA and 5’-TCTTGGCTTCCTATGCTAAATCC; for probe 4 (against IGS1-R, IGS2-R), 5’-TCGCCAACCATTCCATATCT and 5’-CGATGAGGATGATAGTGTGTAAGA and for detecting ACT1, 5’- CGAATTGAGAGTTGCCCCAG and 5’-CAAGGACAAAACGGCTTGGA. Strand-specific probes were then prepared by linear PCR in a final volume of 20 μL containing 0.2 mM dATP, 0.2 mM dTTP, 0.2 mM dGTP, 50 μL [α-^32^P]-dCTP (3,000 Ci/mmol, 10 mCi/ml, Perkin Elmer), 1.25 u ExTaq (TaKaRa), 1x ExTaq buffer, 50 ng PCR product as a template, and 10 μM primer (5’-AGTTCCAGAGAGGCAGCGTA for probe 3, 5’-CATTATGCTCATTGGGTTGC for probe 4). PCR was initiated by a denaturation step at 94°C for 3 min, followed by 35 cycles of amplification (96°C for 20 s, 51°C for 20 s, and 72°C for 30 s), and a final step at 72°C for 3 min. Unincorporated nucleotides were removed using ProbeQuant G-50 Micro Columns (GE Healthcare). The radio-labeled probes were heat-denatured by incubating for 5 min at 100°C, immediately prior to hybridization to the membrane.

The membrane was incubated with 10 mL of ULTRAhyb Ultrasensitive Hybridization Buffer (Thermo Fisher) at 42°C for 1 h. The heat-denatured probe was incubated with the membrane overnight at 42°C. The membrane was rinsed twice with 2x SSC, washed for 15 min at 42°C twice with wash buffer 1 (2x SSC, 0.1% SDS), and washed for 15 min at 42°C twice with wash buffer 2 (0.1x SSC, 0.1% SDS). The membrane was exposed to phosphor screens for several days and radioactive signals were detected using Typhoon FLA 7000 (GE Healthcare). Probes were stripped by incubating the membrane with boiled 0.1% SDS by shaking for ∼30 min, rinsed with 2x SSC, and re-hybridized with the ACT1 probe that was prepared as described above. Signals of IGS-F, IGS1-R, IGS2-R, and ACT1 were quantified using FUJIFILM Multi Gauge version 2.0 software (Fujifilm). The levels of IGS-transcripts were normalized to the ACT-1 signal.

### Replicative lifespan analysis

Replicative lifespan was measured as previously described (Kennedy et al., 1994). In brief, strains were streaked out on a YPD plate at a low density and incubated at 30°C. Cells that emerged as a small bud were placed to other areas of the plate using a Singer micromanipulator system, and the plate was incubated at 30°C. When these cells produced buds, the budded daughter cells were removed using a micromanipulator until the mother cell stopped producing more buds. The number of budded daughter cells was counted and designated as the replicative lifespan of each mother cell. Using replicative lifespan, the survival curve and average life span were determined. Statistical significance was determined by t-test.

### Sorting of old cells

Old cells were sorted as described previously (Sinclair and Guarente, 1997) with modifications. A single colony was inoculated into 5 mL YPD and grown overnight at 30°C. The overnight YPD culture was diluted in 12 mL YPD to an OD_600_ = 0.2 and grown to an OD_600_ = ∼0.7–1.0. Then, 5.0×10^7^ cells were collected by centrifugation for 2 min at 1,800 × *g* at room temperature and washed twice in phosphate-buffered saline (PBS). The cells were resuspended in 200 μL PBS and, after addition of 3.75 mg of sulfo-NHS-LC-biotin (Thermo Fisher Scientific) in 125 μL of PBS, incubated at room temperature with gentle shaking for 15 min. Cells were collected, washed three times in 500 μL PBS, resuspended in 1 ml YPD and split into 500 μL cell suspensions that were used to inoculate two 1L flasks with 250 mL YPD medium. Cells were grown for 12 h at 30°C with shaking, allowing cells to divide ∼8 times. Before OD_600_ exceeded 1.0, cells were collected by centrifugation and resuspended in cold PBS. 100 μL of PBS-washed streptavidin-coated magnetic beads (PerSeptive Biosystems, MA) were added to the cells, followed by incubation at room temperature for 15 min with occasional swirling. Tubes were placed on a magnetic sorter for 5 min at 4°C. The supernatant containing cells unattached to the magnet, was transferred to a new tube and centrifuged. The cell pellet was stored as young cells. The cells attached to the magnet were eluted in cold PBS and washed five times. This cell pellet was stored as old cells. Bud scars were stained with Calcofluor (Sigma), visualized by fluorescent microscopy, and counted (50 cells per sample).

## Supplemental Information

**Figure S1.**
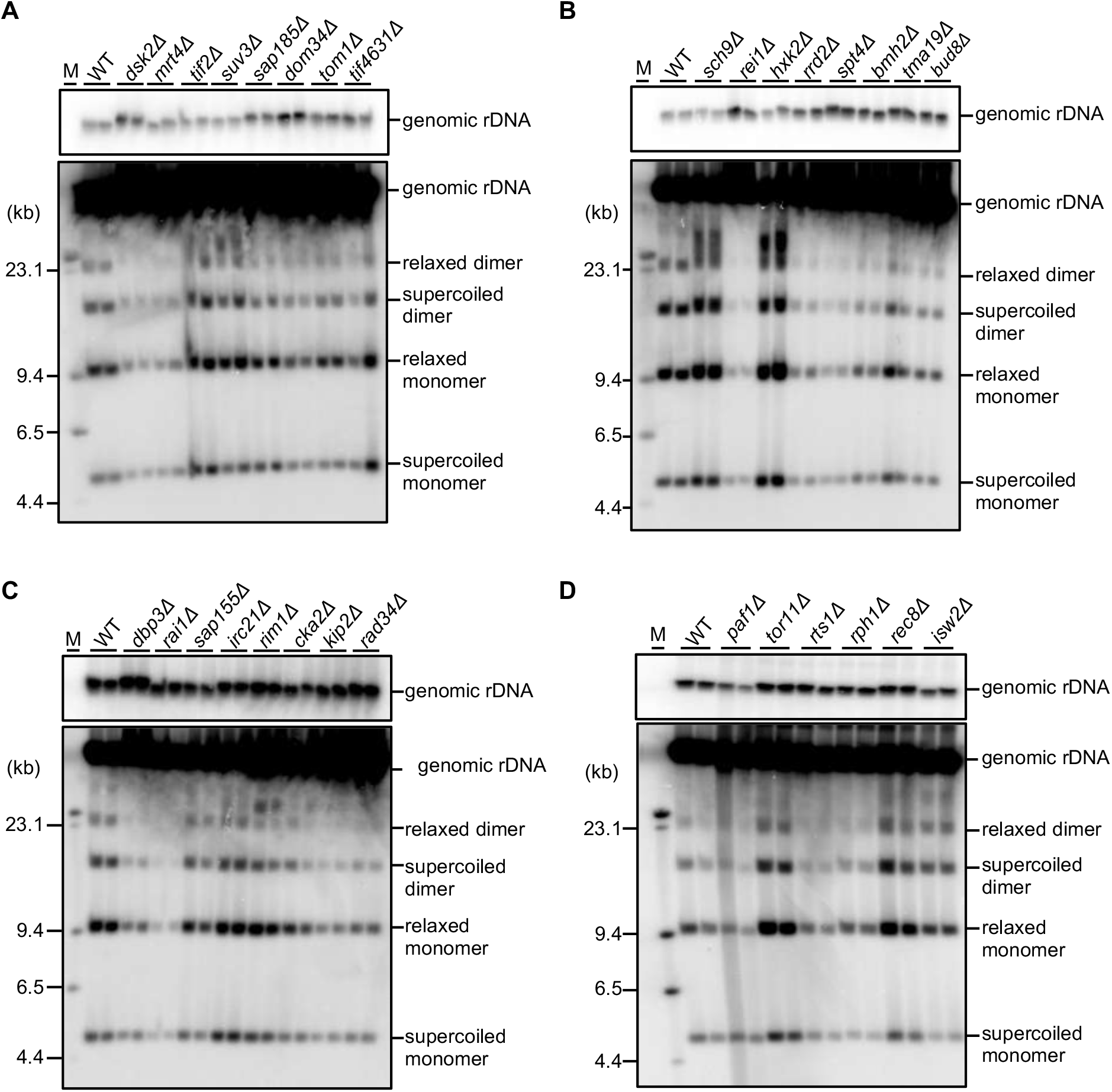
ERC detection. DNA isolated from the indicated mutant strains was separated by agarose gel electrophoresis, followed by Southern blotting with probe 1 (see Fig. 1A). The quantified signal intensities are shown in Fig. 1B. Genomic rDNA, supercoiled and relaxed forms of monomeric and dimeric ERCs are indicated. Sizes of lambda DNA-Hind III markers are indicated.

**Supplementary Table 1.**
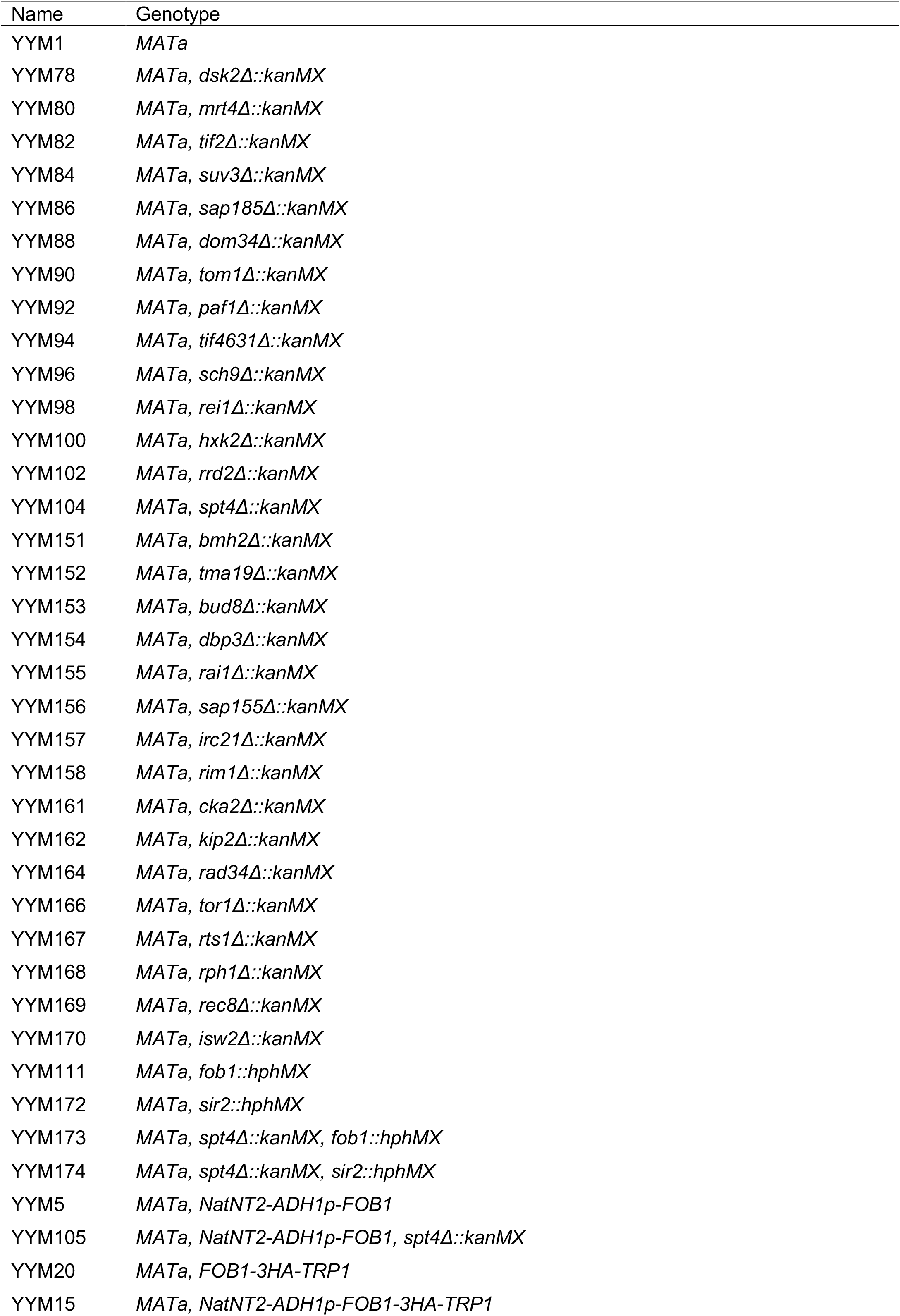
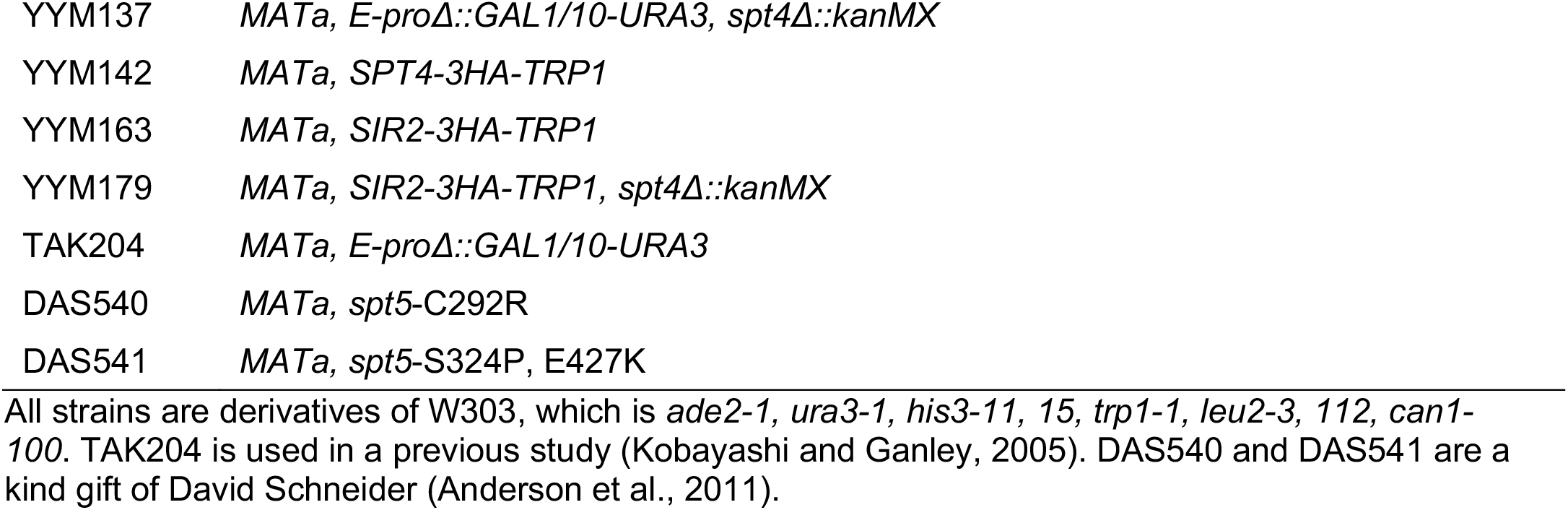
*Saccharomyces cerevisiae* strains used in this study

